# Avobenzone, Guaiazulene and Tioxolone identified as potent autophagy inducers in a high-throughput image based screen for autophagy flux

**DOI:** 10.1101/2022.01.17.476702

**Authors:** Surendra Kumar Prajapat, Chandru Subramani, Puja Sharma, Sudhanshu Vrati, Manjula Kalia

## Abstract

Autophagy is a conserved intracellular degradation pathway that is essential for maintaining cellular homeostasis. Given its critical role in several disease conditions, recent studies are focussed on identifying drugs/small molecules with autophagy modulating capacity for potential clinical applications. Here, we describe the development and characterisation of a quantitative image-based high content screening platform for autophagy flux measurements using the human melanoma A375 cell line that stably expresses the GFP-LC3-RFP probe. The GFP-LC3 is incorporated into autophagosomes, while RFP serves an internal control. The GFP/RFP fluorescence intensity ratio gives an accurate indication of autophagy induction (low ratio) vs blockage of autophagy flux (high ratio), and was validated with the autophagy inducer Torin1 and inhibitor Bafilomycin A_1_. This assay was used to screen the Spectrum collection library comprising of 2560 compounds, to identify autophagy modulators. In addition to known autophagy effectors, several novel autophagy inducers and inhibitors were identified in our study. Further three FDA approved drugs that are widely used in skin-care products: Avobenzone, Guaiazulene and Tioxolone, were validated as potent autophagy inducers that function in an mTOR independent manner.

## Introduction

Macroautophagy (hereafter, autophagy) is an intracellular degradative pathway, conserved from yeast to human. It occurs at a basal level to maintain cellular homeostasis, and is upregulated during starvation and disease conditions [1]. Recent studies have shown that pharmacological modulation of autophagy holds tremendous therapeutic potential for disease conditions such as neurodegeneration and cancer [2], and hence identification of autophagy modulating drugs/compounds is an active area of research [3].

The measurement of autophagy is done by monitoring the lipidated levels of microtubule-associated protein light chain 3 (LC3) protein that specifically incorporates into autophagosomes. Other techniques involve direct visualization of autophagosomes through fluorescence or electron microscopy. The degradative capacity of the pathway or autophagy flux is analysed by using specific inhibitors of autophagosome-lysosome fusion either through western blotting for LC3-II levels or by using ratiometric fluorescence based assays [4–6]. These have been further developed for high throughput platforms and can enable testing the autophagy capacity of several thousand compounds. A series of image based high throughput screening studies using probes such as mRFP-GFP-LC3, GPF-LC3-RFP-LC3ΔG, GFP-LC3-RFP etc. have identified autophagy inducers with broad translation potential [6–10].

Here we have developed an image-based high content quantitative assay using the human melanoma A375 cell line stably expressing the fluorescent probe GFP-LC3-RFP [6]. In a primary screening of the Microsource spectrum library, several autophagy flux inducers and inhibitors were identified by low and high GFP/RFP signal ratios respectively. Three drugs widely used in skin-care products Avobenzone, Guaiazulene and Tioxolone, were further validated for their autophagy inducing properties.

## Results

### Generation and validation of A375 cell line stably expressing autophagy flux probe

Studies have established that the GFP-LC3-RFP probe is a powerful tool for measurements of autophagy flux [5,6,11]. This protein is cleaved by the cellular cysteine protease ATG4, into two equimolar fragments of GFP-LC3 and RFP. The GFP-LC3 is lipidated with phosphatidylethanolamine (PE) at the terminal glycine residue, resulting in its incorporation into autophagosomes. Depending on the status of autophagy in the cell, the levels of GFP-LC3 change, while the RFP serves as an internal control (Fig 1A). The ratiometric measurements of GFP/RFP relative to a basal/untreated condition are an accurate reflection of autophagy induction or inhibition in the cell (Fig 1A) [6,12].

**Figure 1.**
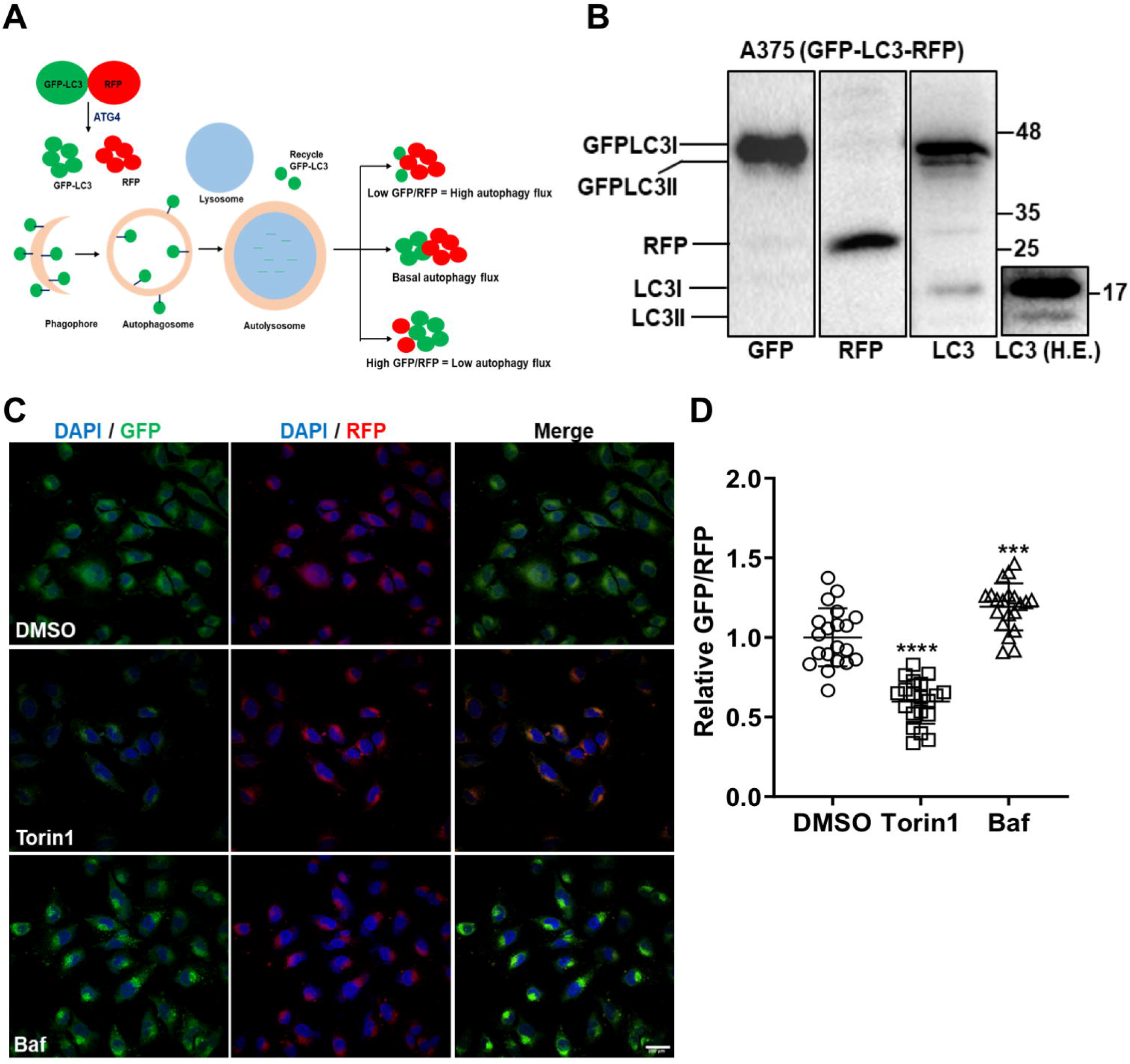
Development of GFP-LC3-RFP cell-line for autophagy flux assays. (A) Schematic representation of the autophagy flux measurement assay using the GFP-LC3-RFP fluorescent probe. (B) The stable A375 (GFP-LC3-RFP) clone was analysed by western blotting using GFP, RFP and LC3 antibodies. Cleavage of the GFP-LC3 from the RFP fragment is observed. Endogenous LC3 bands can be seen in a higher exposure (H.E) blot on the extreme right (C) A375 (GFP-LC3-RFP) cells were treated with DMSO or autophagy inducer Torin1 (1 μM) or inhibitor Baf (100 nM) for 6 h. Cells were imaged on a confocal microscope. Scale bar 200 μm. (D) GFP/RFP intensity ratios were measured from 20-30 cells from two independent coverslips and normalized to DMSO control. One-way ANOVA was used to determine statistical significance, \*\*\**P* < 0.001, **** *P* < 0.0001

We generated a stable human melanoma A375 cell line expressing this autophagy flux probe (Fig 1B). Positive clones were validated, and one clone for chosen for further studies. The GFP-LC3 expression was detected using both GFP and LC3 antibodies (Fig 1B). Roughly equal levels of RFP were also seen (Fig 1B). Fluorescence images showed that the clone expressed GFP-LC3 and showed punctate distribution in the cell corresponding to autophagosomes, while diffuse RFP staining was observed throughout the cytosol (Fig 1C, upper panel). Non-toxic concentrations of Torin1 (autophagy inducer), and Bafilomycin A_1_ (Baf) (autophagosome-lysosome fusion inhibitor), were determined (Fig S1A-B). Treatment of these cells with Torin1, resulted in a clear reduction in GFP-fluorescence in these cells, while treatment with Baf showed enhanced GFP intensity (Fig 1C). GFP/RFP ratios in the Torin1 and Baf treated cells were normalized to DMSO treated cells. Torin1 treatment showed a clear reduction of GFP/RFP ratio indicative of high autophagy flux, while Baf showed an enhanced ratio indicative of low autophagy flux (Fig 1D). These data established the validity of this cell line for autophagy flux measurements.

### Establishment of a high content screening platform

This cell line was next tested in a high throughput 96 well plate platform. Cells were treated with DMSO (control) or Torin1/ Baf, and images were acquired. As seen earlier through confocal microscopy, Torin1 and Baf treatment resulted in reduced and enhanced GFP/RFP ratios respectively (Fig 2A, S1C-D). Based on values obtained with Torin1 and Baf, we established a cut off limit of <0.8 for autophagy inducers and >1.2 for autophagy inhibitors (Fig 2B). Using this technique, we screened the Spectrum library of 2560 compounds at 10 μM concentration. Control, Torin1, Baf, and drug treatments were given for 12 h. The cytotoxicity of the drug treatment was established through DAPI staining compared to untreated controls as described in the methods section (Data Sheet S1). Based on the GFP/RFP ratios, we identified 104 drugs that changed the cellular autophagy levels (Fig 2C-D, Fig S2, File S1). The autophagy inducers (Fig 2C) and inhibitors (Fig 2D) were classified based on their reported biological activities in literature. Of these 53 drugs have been previously established as autophagy inducers in literature [6,7,9]. We further identified 31 novel autophagy inducers and 19 novel autophagy inhibitors (File S1).

**Figure 2.**
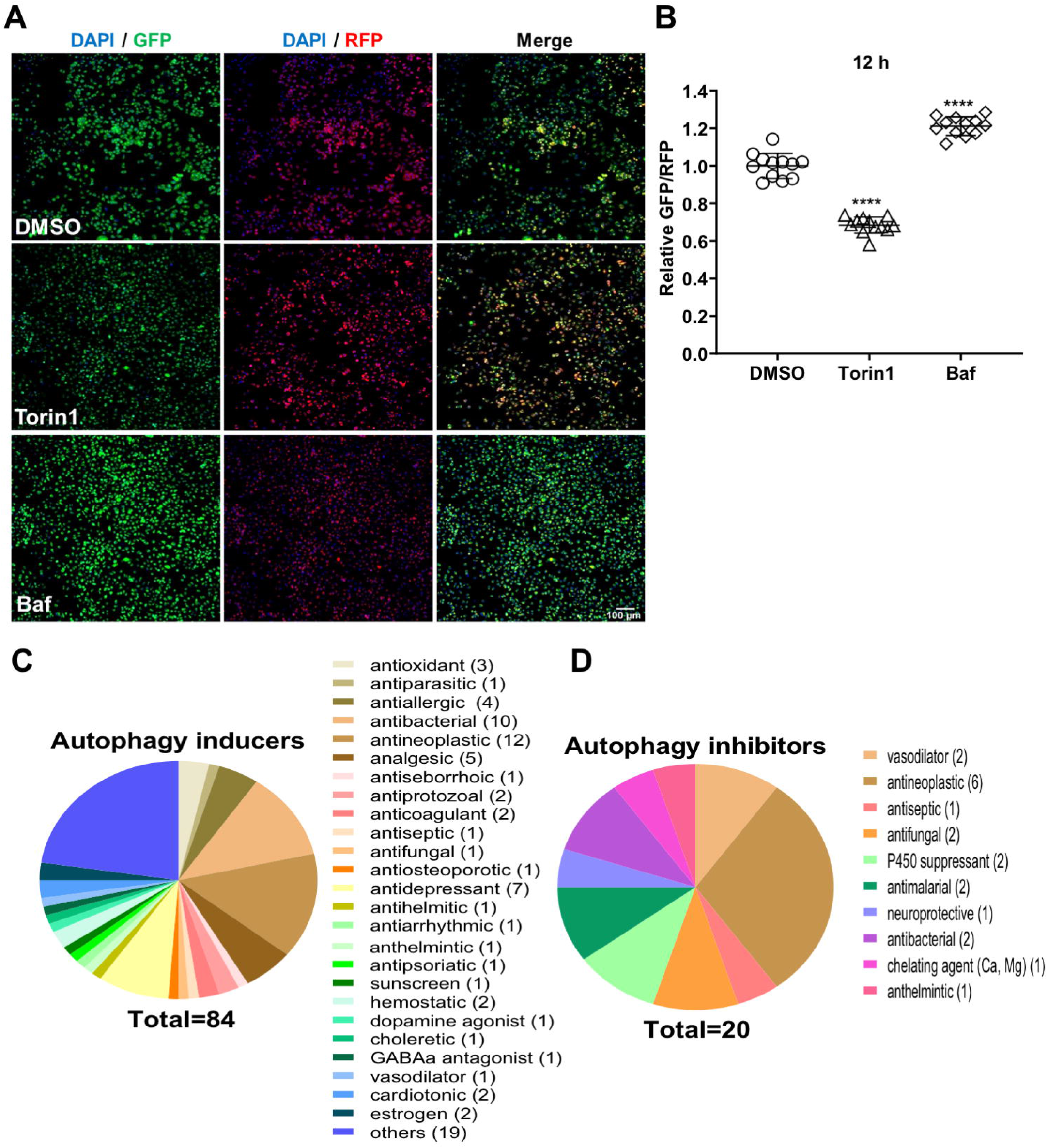
High throughput imaging screen of Microsource spectrum library. (A) The Microsource spectrum library consisting of 2560 drugs were screened for the identification of autophagy flux modulators. A375 (GFP-LC3-RFP) cells were treated with DMSO/ Torin1 (100 nM)/ Baf (20 nM)/ drugs (10 μM) for 12 h in biological duplicates or triplicates. Images were acquired using the Image Xpress high content imaging system, and representative images are shown. Scale bar 100 μm. (B) GFP/RFP ratios were measured and normalized to DMSO control. Values show relative GFP/RFP ratios from 3 independent experiments. (C-D) Venn diagram showing the categorical classification of different molecules identified as autophagy inducers (ratio <0.8) or inhibitors (ratio >1.2). One-way ANOVA was used to determine statistical significance, **** *P* < 0.0001

### Avobenzone, Guaiazulene and Tioxolone identified as potent autophagy inducers

Of the shortlisted drugs, we focused our attention on three FDA-approved drugs that are widely used in skin-care products: Avobenzone, Guaiazulene and Tioxolone. All the drugs were retested for toxicity (Fig 3A), and showed GFP/RFP ratios comparable to Torin1 (Fig 3B, C), identifying them as potent autophagy inducers. We also checked these drugs with the ptfLC3 reporter [5], and observed an increase in the number of autophagosomes (mRFP^+^/GFP^+^) and autolysosomes (mRFP^+^/GFP^-^) on drug treatment, that was comparable to Torin1 (Fig S3). Lysosome acidification was checked with the LysoSensor Yellow-Blue assay [13], and was found to be similar to Torin1 treatment (Fig S4). We further validated autophagy induction through western blotting for LC3. All the drugs showed enhanced LC3-II levels (Fig 4A-C). By using Baf we confirmed that LC3-II accumulation caused by these drugs was due to autophagy induction and not due to blockage of autophagosome-lysosome fusion (Fig 4A-C). Levels of p62 clearance were also assessed through western blotting and these showed a reduction compared to DMSO treatment (Fig 4D-E), but did not have any effect on p62 mRNA levels (Fig 4F). We also tested if these drugs acted in an mTOR dependent manner by checking phosphorylation levels of S6 kinase (S6K) and eukaryotic translation initiation factor 4E-binding protein 1 (4E-BP1). While Torin1 resulted in efficient dephosphorylation of S6K and 4E-BP1, these drugs had no effect on phosphorylation of S6K and 4E-BP1 (Fig 4G-H), suggesting that their mode of action is mTOR independent.

**Figure 3.**
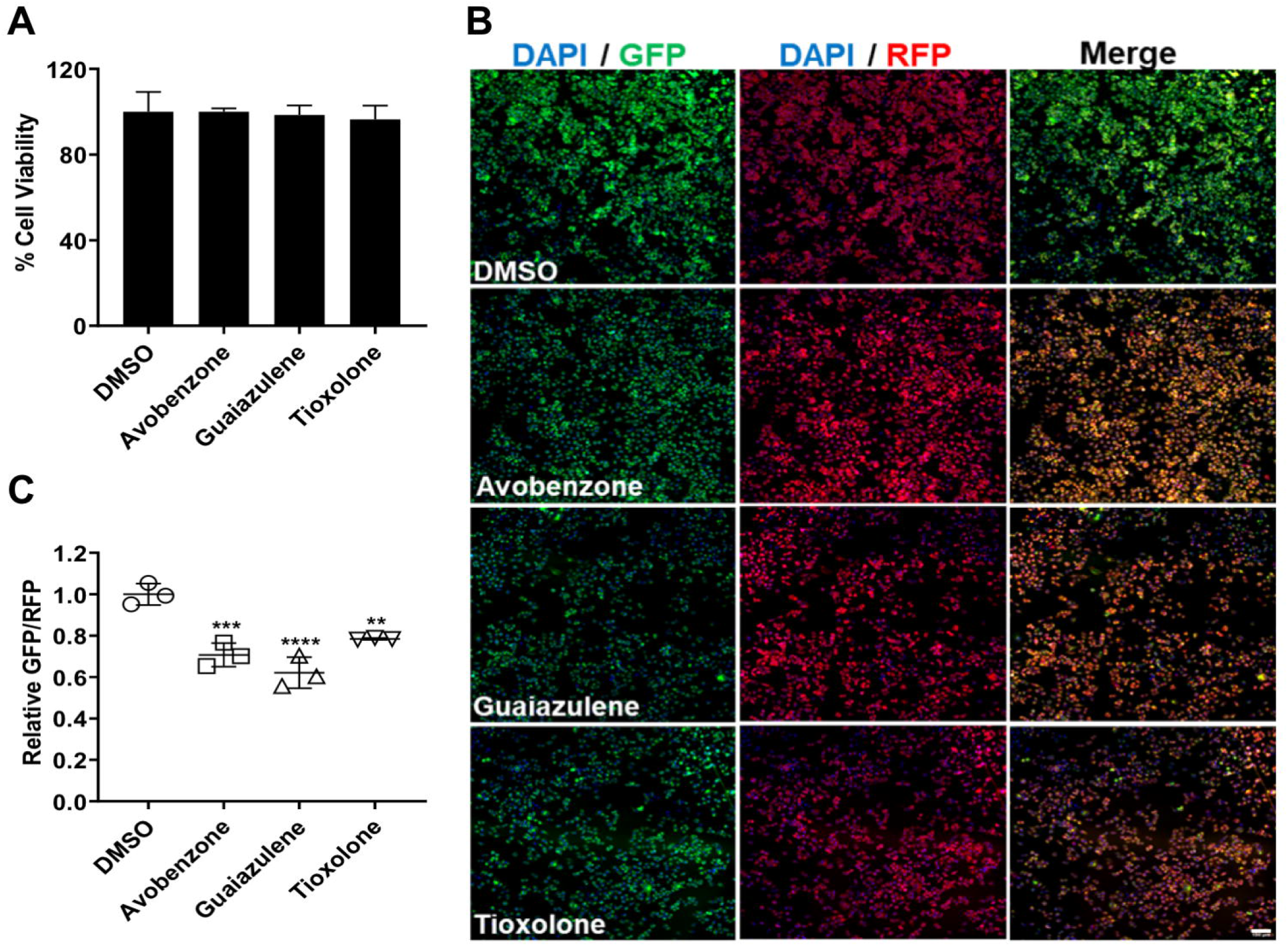
Avobenzone, Guaiazulene and Tioxolone are autophagy inducers. (A) Viability measurements of A375 cells treated with Avobenzone/ Guaiazulene/ Tioxolone (10 μM) for 12 h. (B-C) A375 (GFP-LC3-RFP) cells were treated with DMSO, drugs (Torin1/ Baf) or Avobenzone/ Guaiazulene/ Tioxolone (n=3), and images were acquired using the Image Xpress high content imaging system. The relative ratiometric quantitation of GFP/RFP normalized to DMSO control is plotted in (C). One-way ANOVA was used to determine statistical significance, \*\*\**P* < 0.001, **** *P* < 0.0001.

**Figure 4.**
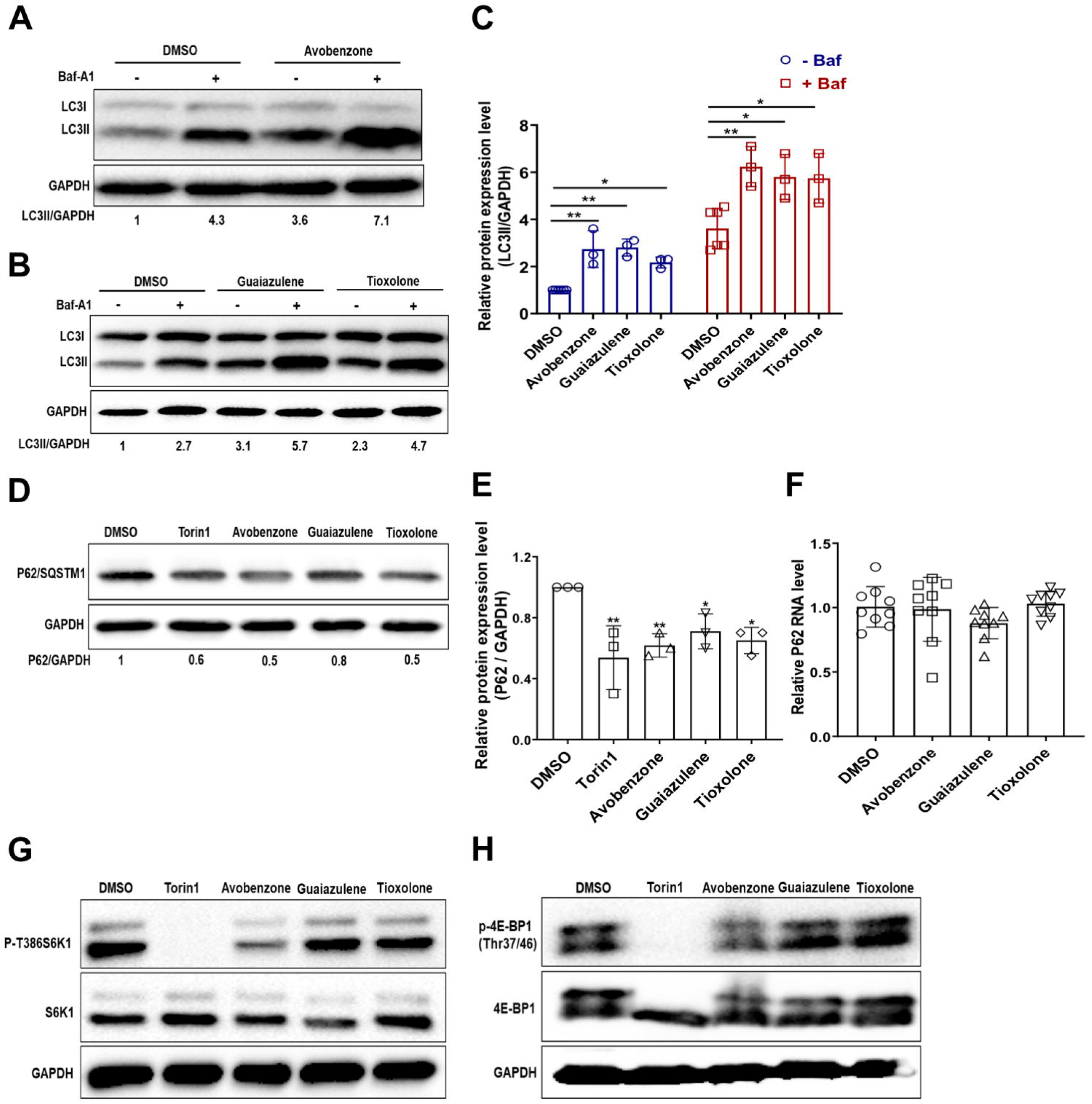
Characterization of Avobenzone, Guaiazulene and Tioxolone as autophagy inducers. (A-B) A375 cells were treated with the indicated drugs (10 μM) for 10 h, followed by Baf treatment (100 nM) for 2 h before harvest and immunoblotted with LC3 and GAPDH antibodies. Increase in LC3-II levels on Baf treatment is indicative of functional autophagy flux. Numbers below the blot indicate ratio of LC3-II/GAPDH, calculated using Image J. (C) Bar graph shows relative quantitation of LC3 expression in Baf treated cells normalized to GAPDH from three independent experiments. (D-E) Cells were treated as described for panel A, and lysates were blotted for p62 and GAPDH. (E) Bar graph shows quantitation of p62 levels normalized to GAPDH from three independent experiments. (F) Relative mRNA levels of p62 in cells treated as described above. (G-H) mTOR downstream signaling was checked by measuring the proteins level of p-T386S6K1, S6K1, p-4E-BP1(Thr37/46) and 4E-BP1 antibodies. Torin1 treatment was used as a positive control. Data is representative of three independent experiments. One-way ANOVA was used to determine statistical significance, \**P* < 0.05, \*\**P* < 0.01, \*\*\**P* < 0.001.

## Discussion

Autophagy research has rapidly evolved and interest has heightened on examining the potential of novel/existing drugs to modulate autophagy [14]. Autophagy modulation is likely to have therapeutic benefits in several disease conditions [15]. Conversely, several FDA-approved drugs could be autophagy modulators but have not been studied from this perspective.

In our current study, we developed a reporter cell line for high throughput measurements of autophagy flux, and identified several known and novel autophagy modulators in the Spectrum collection library of 2560 compounds. Our primary screening and validation showed that three FDA-approved drugs that are widely used in skin-care preparations: avobenzone, guaiazulene, and tioxolone can efficiently induce autophagy flux in an mTOR independent manner.

Avobenzone (butyl methoxy dibenzoylmethane; parsol 1789) is an ultraviolet A absorber widely used in sunscreens [16–22]. Studies have shown that avobenzone can efficiently penetrate the stratum corneum and viable epidermis [23]. It has been shown to protect against UVA-induced melanogenesis through indirect regulatory effect on the Nrf2-ARE pathway [24]. It has also been shown to inhibit the proliferation of human trophoblasts, enhance Akt and ERK1/2 activity and induce mitochondrial membrane depolarization [25]. In normal human epidermal keratinocytes, it can downmodulate lipid metabolism, and Peroxisome proliferator-activated receptor-gamma signaling [26], and can induce keratinocyte derived Vascular endothelial growth factor production [27]. Its autophagy inducing properties have not been described till date, however based on the above-mentioned studies, it is likely that autophagy will play an important role in the mode of action of avobenzone.

Guaiazulene (1,4-dimethyl-7-isopropylazulene), is a natural azulenic compound widely used in cosmetic and health-care products and in pharmaceutical preparations such as creams and toothpastes [28,29]. Its early use was as an ophtlamic drug, and later it became a popular skin conditioning agent. The drug was also shown to be promising for the treatment of gastritis and peptic ulcers [30]. Studies have shown that it can act in a synergic manner with diclofenac and can have therapeutic advantages for the clinical treatment of inflammatory pain [31]. It has also been shown to have anti-cancer properties [32]. A recent study showed that guaiazulene inhibited Akt/mTOR signalling and induced autophagosome formation [33]. However, this study used a very high concentration of guaiazulene (100/150 μM), which might account for the observed mTOR inhibition.

Tioxolone (6-Hydroxy-2H-1,3-benzoxathiol-2-one) is used for cosmetics products such as hair shampoos, skin cleansers and acne treatment products [34]. It has anti-fungal, anti-bacterial, anti-inflammatory and anti-tumorigenic effects, and has been shown to be an inhibitor of human carbonic anhydrase II [35,36]. Tincture of thioxolone plus benzoxonium chloride was shown to be useful for the treatment of cutaneous leishmaniasis [37].

In summary, our high throughput screening approach has identified several novel autophagy inducers and inhibitors. We also validate that three widely used drugs in skin-care products are potent autophagy inducers. Further studies will highlight if the autophagy inducing properties of these drugs govern their mode of action.

## Materials and Methods

### Cell lines

A375 and HEK293T cells were obtained from the Cell Repository at the National Centre for Cell Sciences, Pune, India. Cells were grown in Dulbecco’s modified Eagle’s medium (DMEM) (HiMedia, AL007A) supplemented with 10% fetal bovine serum (FBS) (HiMedia, RM10432), 100 μg/ml penicillin-streptomycin, 2 mM L-glutamine and 1x MEM Non-Essential Amino Acids Solution (HiMedia, ACL006).

### Reagents, antibodies and plasmids

The following reagents were used in the study: Torin1 (Tocris Bioscience, 4247), Bafilomycin (Baf)-A1 (Sigma-Aldrich, B1793-10UG), PMSF (Sigma-Aldrich, P7626-100G), Protease inhibitor cocktail (Sigma-Aldrich, 11697498001), LysoSensor Yellow/Blue DND-160 (Thermo Fisher Scientific, L7545). The Spectrum collection library (MicroSource Discovery Systems, Inc., SP170615) comprising of 2560 compounds including bioactive molecules, natural products, and FDA-approved compounds was used for image based high content screening. The following antibodies were used in the study: GAPDH (14C10); Phospho-4E-BP (Thr37/46) (2855S); 4E-BP (9644S); Phospho-p70 S6 Kinase (Thr389) (97596S) and p70 S6 Kinase (2708S) from Cell Signalling Technology, LC3 (ab51520); RFP (ab62341), GFP (ab32146) and SQSTM1/p62 (ab56416) from Abcam, HRP-conjugated secondary antibodies from Jackson ImmunoResearch Laboratories. The following plasmids were obtained from Addgene: gag/pol (14887, Tannishtha Reya) [38], pCI-VSVG (1733, Garry Nolan) [39], pMRX-IP-GFP-LC3-RFP (84573, Noboru Mizushima) [6], and ptfLC3 (21074, Tamotsu Yoshimori) [10]. The following primers (5□-3□) were used in the study GAPDH: F-TGCACCACCAACTGCTTAGC, R-GGCATGGACTGTGGTCATGAG; P62/SQSTM1: F-AGGCGCACTACCGCGAT, R-CGTCACTGGAAAAGGCAACC.

### Stable cell line generation

HEK293T cells were co-transfected with gag/pol, pCI-VSVG and pMRX-IP-GFP-LC3-RFP plasmids using Lipofectamine 2000 and incubated at 37°C for 24 h. The supernatant was harvested for retrovirus, and was used to transduce A375 cells. Puromycin selection (2 μg/ml) was given after 48 h. After one week of selection, cells were trypsinized and 10,000 cells were serially diluted to achieve single cell seeding per well of a 96 well-plate. Isolated colonies were visible by 14 days, and were tested for the expression of GFP, RFP and autophagy flux using Torin1 and Baf treatments. One clone was chosen for further studies.

### Drug treatment & confocal imaging

Stable GFP-LC3-RFP A375 cells or A375 cells transiently transfected with ptfLC3, were treated with Torin1 (1 μM/ 6h; 100 nM/ 12h) or Baf (100 nM/ 6h; 20 nM/ 12h) or DMSO/Avobenzone/ Guaiazulene Tioxolone (10 μM) for 12 h. For confocal microscopy all cells were grown on glass coverslips. GFP/RFP intensity ratio per cell was calculated and normalized to DMSO control treated cells. For ptfLC3 experiments, autophagosomes (RFP^+^/GFP^+^ structures) and autolysosomes (RFP^+^/GFP^-^ structures) per cell were counted using Spot to Spot colocalization analysis in Imaris 8.

### High content imaging and analysis

For image based high content screening 15,000 cells/well were seeded in Corning 96 well black polystyrene microplates (Corning, CLS3603). Torin1 (100 nM), Baf (20 nM), and drug (10 μM) treatment was given for 12 h. All drugs were tested in biological duplicates or triplicates. Cells were fixed and stained with DAPI. Images were acquired on ImageXpress Micro Confocal High-Content Imaging System (Molecular Devices, USA) using FITC, Texas red, and DAPI channels with a 10 X objective lens. A total of 16 fields per well were acquired and this covered the entire well area. Analysis was performed using the multiwavelength cell scoring module of the MetaXpress Software. To calculate drug cytotoxicity, total nuclei were counted using DAPI in drug and DMSO treated wells. Any drug showing more than 20% reduction in total nuclei was considered as toxic and excluded from further analysis. Only triple positive cells (DAPI, GFP, RFP) were used for quantitation. The integral intensity of GFP and RFP per well was calculated and used to estimate the GFP/RFP ratio. The values obtained for DMSO treatment were used for normalization. The GFP/RFP ratio in Torin1 treatment gave a mean value of 0.68, while BafA1 treatment gave a mean value of 1.2. Any drug showing a GFP/RFP ratio <0.8 was considered as an autophagy inducer, and GFP/RFP ratio >1.2 was marked as autophagy flux inhibitor.

### Western blotting

After drug treatment A375 cells were washed with PBS and lysed using cell lysis buffer [150 mM NaCl, 1% Triton X-100, 50 mM Tris-HCl (pH 7.5), 200 μM PMSF and protease inhibitor cocktail]. Protein quantification was done using BCA assay (G-Bioscience, 786-570) kit. Cell lysates were separated on polyacrylamide gels and transferred to a PVDF membrane for immunoblotting using specific primary and secondary antibodies. Western blot images were processed using ImageJ (NIH, USA) software for quantitation of band intensities. Data are represented as mean ± SD from three independent experiments.

### LysoSensor Yellow-Blue assay

The LysoSensor Yellow-Blue assay was performed as described in [13]. A375 cells were were grown on glass coverslips, and were treated with DMSO (control), Torin1 (1 μM) or Baf (100 nM) or Avobenzone/ Guaiazulene Tioxolone, followed by incubation with 10 μM LysoSensor for 5 min. Cells were then washed with ice-cold PBS three times and fixed using 4% PFA. Cells were imaged on LSM 880, Carl Zeiss, with excitation wavelength range 371-405 nm and emission wavelength range 420-650 nm. The LysoSensor dye has dual-emission peaks of 440 nm (blue in less acidic organelles) and 540 nm (yellow in more acidic organelles). Yellow fluorescence per cell was quantified and normalized to DMSO treated control.

### Cell viability assay

A375 cells were seeded at a density of 15000 cells/well in 96-well plates. After 24 h, cells were treated with indicated concentrations of Torin1/ Baf/ Avobenzone/ Guaiazulene Tioxolone for 6 or 12 h. The cytotoxicity assay was performed using CellTiter-Glo kit (Promega, G7572). Percentage of cell viability was measured relative to DMSO treated cells.

### Statistical analysis

Statistical analysis of the data was performed using one-way ANOVA and differences were considered significant at values of \**P* < 0.05, \*\**P* < 0.01, \*\*\**P* < 0.001, **** *P* < 0.0001

## Abbreviations

Baf: Bafilomycin A_1_
LC3: Microtubule-associated protein light chain 3
mTOR: mechanistic target of rapamycin

## Acknowledgements

High content screening was performed at the Advanced Technology Platform Centre (ATPC) at RCB. We are thankful to Dr. Nirpendra Singh, Ashish Pandey and Rajan for technical support. Simran Chhabra and all other Virology lab members are acknowledged for their support and suggestions.

