## Supplementary Data for "Avobenzone, Guaiazulene and Tioxolone identified as potent autophagy inducers in a high-throughput image based screen for autophagy flux"

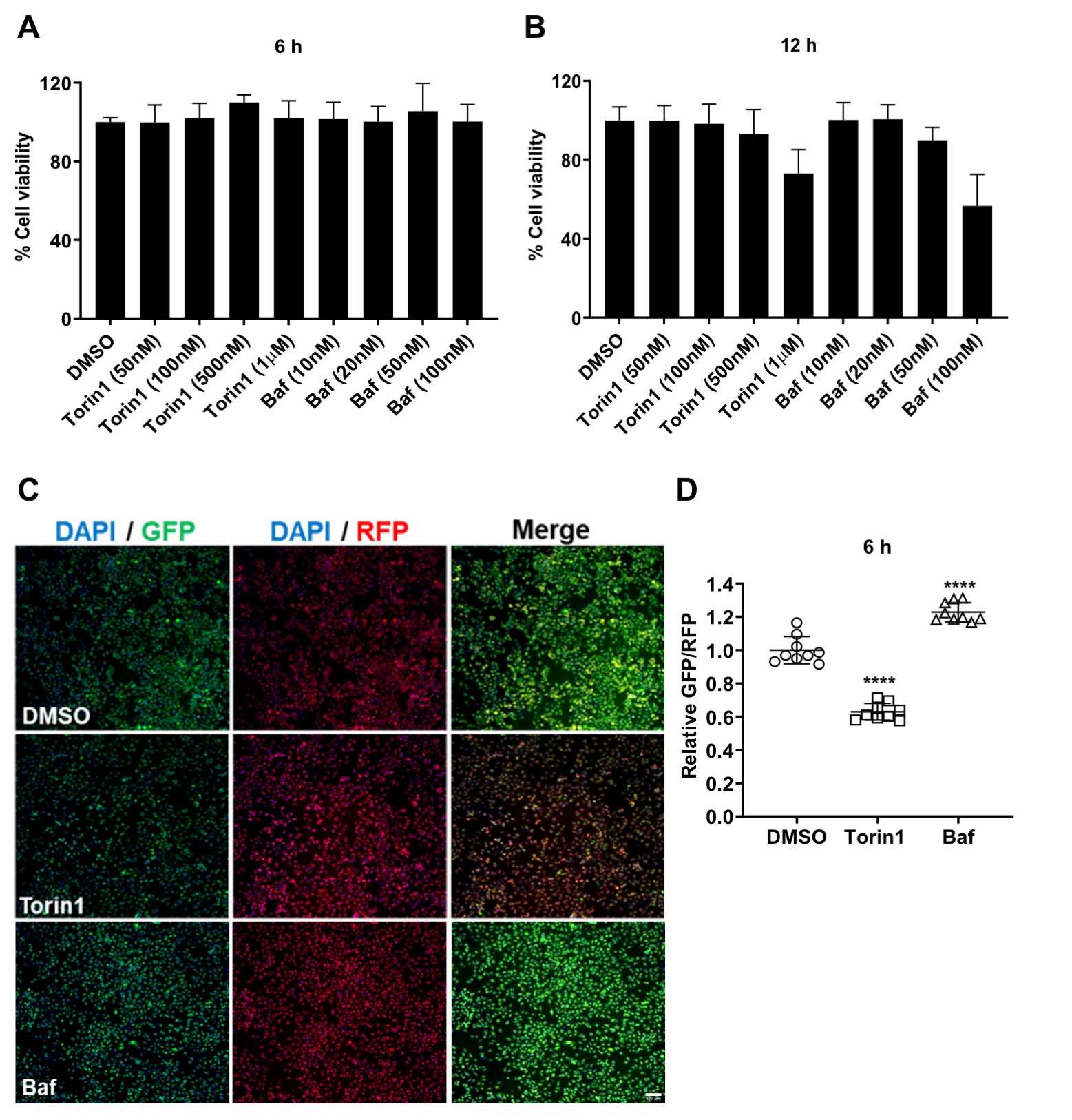

**Fig S1: Establishment of controls for high throughput imaging.** (A-B) Viability measurements of A375 cells treated with indicated concentrations of Torin1/Baf for 6 and 12 h. (C-D) A375 (GFP-LC3-RFP) cells were treated with Torin1 (1 µM), Baf (100 nM) for 6 h. Images were acquired using the Image Xpress high content imaging system, and representative images are shown. Scale bar 100 µm. (D) Values show relative GFP/RFP ratios from 9 independent experiments. One-way ANOVA was used to determine statistical significance, **** *P* < 0.0001

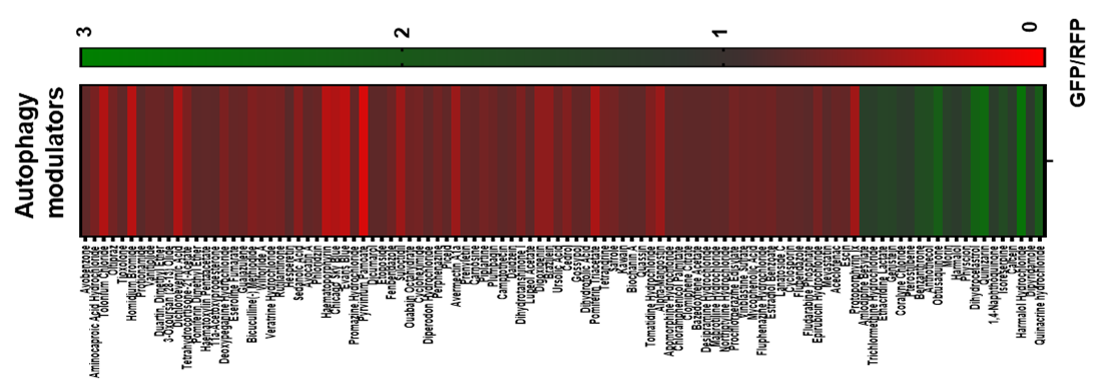

**Figure S2:** Clustered heatmap showing GFP/RFP ratios of autophagy modulators identified in the screen.

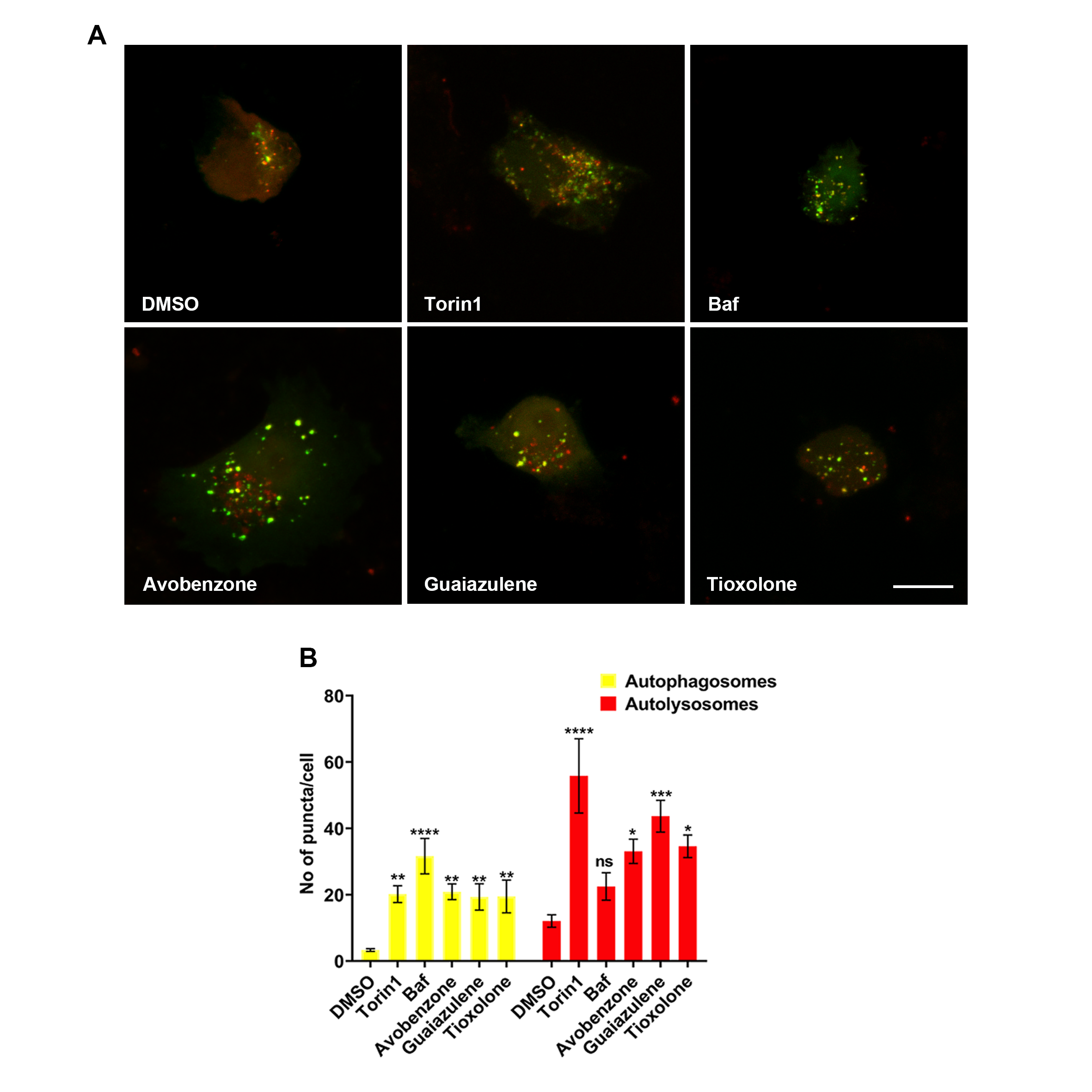

Fig S3: **Avobenzone, Guaiazulene and Tioxolone induce functional autophagy flux.** (A-B) A375 cells were transfected with ptfLC3 and were treated with DMSO, Torin1 (100 nM), Baf (20 nM) or the indicated drugs (10 µM) for 12 h. Cells were imaged on Leica SP8 confocal microscope. Scale bar 10 µm. (B) Bar graph showing number of autophagosomes (RFP^+^/GFP^+^) and autolysosomes (RFP^+^/GFP^−^) calculated from 10-14 cells from two independent experiments plotted as mean ± SEM. Statistical analysis was performed using one-way ANOVA, **P* < 0.05, ***P* < 0.01, ****P* < 0.001, **** *P* < 0.0001 **** *P* < 0.0001.

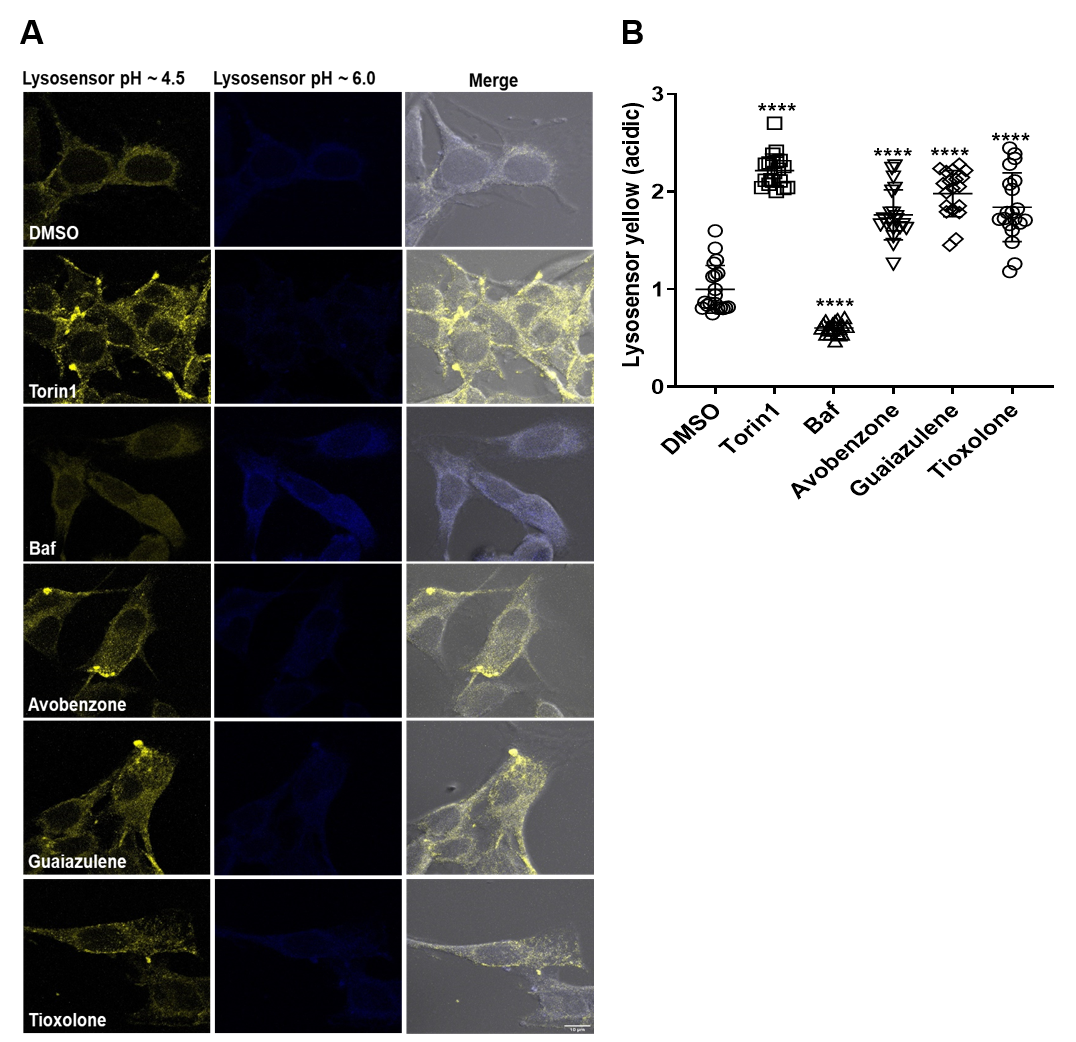

**Figure S4: Avobenzone, Guaiazulene and Tioxolone do not alter lysosome acidification** (A-B) A375 cells were treated with Torin1 (1 µM), Baf (100 nM) for 6 h, or with the DMSO/drugs (10 µM) for 12 h, followed by incubation with 10 µM LysoSensor for 5 min. (B) Yellow fluorescence was quantified from 20 cells from two independent coverslips and normalized to DMSO control. Statistical analysis of the data was performed using one-way ANOVA, **** *P* < 0.0001

**File S1:** Autophagy flux primary screen data showing % nuclei count and GFP/RFP ratios for the 2560 compounds of the Microsource spectrum library (Sheet: GFP,RFP ratios); list of novel and known autophagy inducers (Sheet: inducers); list of novel and known autophagy flux inhibitors identified in the study (Sheet: inhibitors). PMID’s for the known autophagy inducers and inhibitor are indicated.
